# Functional ultrasound imaging reveals pathway-specific visual system reorganization in young *Cln3*−/− mice

**DOI:** 10.64898/2026.06.04.730118

**Authors:** Fanchao Yin, Yanya Ding, Hayley Chang, Viollandi Prifti, Jingyu Feng, Edward G Freedman, John J Foxe, Marvin Doyley, Kuan Hong Wang

## Abstract

CLN3 disease, or juvenile Batten disease, is a neurodegenerative lysosomal storage disorder in which visual impairment is typically the earliest clinical manifestation. Although retinal pathology has been extensively studied, functional alterations within central visual pathways remain poorly understood. Here, we used functional ultrasound (fUS) imaging to characterize visually evoked activity across central visual circuits in young Cln3 knockout (*Cln3−/−)* mice before the onset of severe retinal degeneration. Visually evoked hemodynamic responses were quantified in regions spanning the geniculostriate and extrageniculate visual pathways, including cortical, thalamic, and midbrain regions. To assess regional pathological burden, accumulation of subunit c of mitochondrial ATP synthase (SCMAS), a pathological marker of CLN3 disease, was examined using immunohistochemistry. We found that *Cln3−/−* mice exhibited pathway-specific alterations in visually evoked activity. Regions along the extrageniculate pathway, including the midbrain, posterior thalamus, and anterior secondary visual cortex, showed enhanced activation relative to wild-type controls. In contrast, activation within the geniculostriate pathway was reduced in the anterior thalamus and remained unchanged in the primary and posterior secondary visual cortex. SCMAS accumulation was elevated across all examined visual regions in *Cln3−/−* mice relative to wild-type controls, with greater accumulation observed in geniculostriate regions than in extrageniculate regions. These findings demonstrate early pathway-specific functional and pathological alterations in the visual system of *Cln3−/−* mice, suggesting pathway-level reorganization of central visual processing. This study advances understanding of central visual dysfunction in CLN3 disease and highlights fUS imaging as a sensitive approach for detecting early functional abnormalities in neurodegenerative disorders.

## Introduction

CLN3 disease (Batten disease), the juvenile form of neuronal ceroid lipofuscinosis (NCL), is a rare lysosomal storage disorder (LSD) marked by progressive accumulation of storage materials within neuronal lysosomes (Adams et al., 2013; Consortium, 1995; Munroe et al., 1997). Visual deficits are typically the first clinical manifestation, followed by cognitive decline, motor dysfunction, epileptic activities, and language impairment (Mink et al., 2013; Morison et al., 2025; Ostergaard, 2016). Affected individuals aged 4 to 8 years typically exhibit progressive loss of visual acuity, accompanied by photophobia, impaired color discrimination, and reduced contrast sensitivity (Kuper et al., 2021; Ouseph et al., 2016). Because visual symptoms emerge early and progress rapidly, characterizing functional responses along the visual pathways is essential for understanding disease mechanisms and for developing and evaluating targeted therapeutic strategies.

The neural architecture of visual processing provides a mechanistic framework for interpreting the visual deficits observed in CLN3 disease. The geniculostriate pathway, through which retinal ganglion cells (RGCs) project to the lateral geniculate nucleus (LGN) of the thalamus and subsequently to the primary visual cortex (V1) and higher-order visual areas (Dhande et al., 2015; Garey & de Courten, 1983), supports the conscious perception of form, color, and spatial information (Conway, 2009; Lennie & Movshon, 2005). In parallel, the extrageniculate pathway comprises retinal projections to the superior colliculus (SC) and pretectal nuclei (PN), with relays through the pulvinar/lateral posterior (LP) thalamus and subsequent projections to higher-order visual and associative cortices (Bennett et al., 2019; Bridge et al., 2016). This pathway supports visuomotor coordination, visuospatial orientation, and reflexive visual behaviors (Ahmadlou et al., 2018; Han & Bonin, 2024; Mundinano et al., 2018). While clinical manifestations of CLN3 disease are often studied in terms of ocular pathology (Ouseph et al., 2016; Wright et al., 2020), the contribution of central visual pathways to disease-related visual dysfunction remains poorly understood.

Although lysosomal storage accumulation and neuronal loss have been identified in cortical and subcortical brain regions in both CLN3 human tissue and animal models (Bozorg et al., 2009; Seigel et al., 2002; Weimer et al., 2006), it remains unclear whether functional abnormalities in central visual pathways emerge prior to overt neurodegeneration. This question is particularly important because retinal function in young *Cln3* mouse models appears relatively preserved during early disease stages, with electroretinogram deficits becoming more pronounced only at later ages (around one year old)(Seigel et al., 2002; Volz et al., 2014). In other neurodegenerative and ophthalmic disorders, functional imaging studies have shown that abnormalities within the LGN and V1 are associated with impaired color discrimination and contrast sensitivity (Mullen et al., 2008; Powell et al., 2025). Given that CLN3 disease presents with similar visual deficits (Mink et al., 2013; Morison et al., 2025; Ostergaard, 2016), assessing visually evoked activity within central visual pathways may reveal early circuit-level dysfunction associated with disease progression. Moreover, migraine studies implicate the extrageniculate pathway in photophobia generation and light-induced headaches (Maleki et al., 2012; Noseda et al., 2019). Importantly, photophobia has been reported as an early clinical feature of CLN3 disease and is often a primary complaint during ophthalmologic evaluation (Kuper et al., 2021; Mink et al., 2026). Together, these observations motivate a pathway-level analysis of both geniculostriate and extrageniculate visual circuits in early-stage *Cln3−/−* mice. Achieving this goal requires an imaging modality capable of assessing functional activity across distributed visual networks.

Functional ultrasound (fUS) imaging is a neuroimaging technique that measures changes in cerebral blood flow with high spatial and temporal resolution. Since its development and validation (Mace et al., 2011; Mace et al., 2013), fUS has become an effective tool for mapping brain-wide functional activity. Its physiological basis is grounded in neurovascular coupling, which links neuronal firing to local blood flow and has been validated across multiple modalities. Early studies combining fUS with EEG demonstrated a close correspondence between fUS signals and electrophysiological activity in freely moving rats (Sieu et al., 2015). Subsequent studies combining two-photon microscopy with fUS further linked changes in cerebral blood volume (CBV) to neuronal calcium activity (Aydin et al., 2020), while simultaneous intracranial electrophysiology and fUS recordings in awake mice confirmed tight coupling between hemodynamic and neuronal signals (Nunez-Elizalde et al., 2022). Beyond its physiological validity, fUS offers several practical advantages for functional brain mapping, including high spatiotemporal resolution and sufficient penetration to access deep brain structures. By directly measuring blood flow in the cerebral microvasculature with a spatial resolution of approximately 100 *µ*m and a penetration depth exceeding 10 mm (Deffieux et al., 2018; Montaldo et al., 2022), fUS enables simultaneous imaging of cortical and subcortical regions in the rodent brain. Compared to functional magnetic resonance imaging (fMRI), which relies on blood-oxygen-level–dependent (BOLD) signals and typically offers millimeter-scale spatial resolution in standard settings (Glover, 2011; Logothetis, 2008; Olman & Yacoub, 2011; Ugurbil, 2021), fUS provides greater spatial detail in small-animal imaging. It also achieves greater penetration depth than optical imaging approaches, which provide high spatial resolution but are typically limited to superficial cortical layers (Chhetri et al., 2015; Devor et al., 2012; Ferrari & Quaresima, 2012; Soekadar et al., 2021). Together, these characteristics place fUS in a favorable position to bridge the gap between macroscopic whole-brain techniques and high-resolution optical approaches (Robin et al., 2021), making it particularly well-suited for investigating multiple levels of brain organization along visual processing pathways.

The feasibility and reliability of fUS for investigating the visual system have been clearly demonstrated in rodents, particularly in mice and rats. Early work in mice showed that whole-brain fUS can detect widespread visual and visuomotor activation during optokinetic stimulation, identifying dozens of active brain regions and revealing reflex-dependent functional modules (Mace et al., 2018). Additional studies demonstrated volumetric fUS imaging of brain activity in awake mice during visual tasks and optogenetic stimulation (Brunner et al., 2020), followed by the introduction of a framework for whole-brain fUS imaging that established the practicality of mapping visual responses at high spatiotemporal resolution (Brunner et al., 2021). More recently, fUS has been used to characterize region-specific hemodynamic profiles across the mouse visual system (Erol et al., 2023), supporting its ability to resolve spatially differentiated responses. In rats, fUS combined with visual stimulation has enabled accurate mapping of activation across different regions under anesthesia (Gesnik et al., 2017). In spontaneously hypertensive rats, fUS revealed increased visual response intensity in the visual cortex and superior colliculus (Droguerre et al., 2022). Beyond rodent models, fUS has also been successfully applied to study visual processing in other species, including ferrets, where it revealed fine-scale functional organization across the full cortical depth of visual cortex and thalamus during visual stimulation (Hu et al., 2023), and non-human primates, where it enabled imaging of visually driven activity in deep visual and prefrontal regions during both rest and visually guided behaviors (Blaize et al., 2020; Dizeux et al., 2019). Collectively, these studies demonstrate the robustness and versatility of fUS for characterizing neural dynamics within the visual system across species and experimental paradigms, establishing a strong foundation for its application to disease models.

In the present study, we used functional ultrasound imaging (fUS) to examine visually evoked responses across parallel central visual pathways in young adult (3-month-old) *Cln3−/−* mice. We observed increased activation in regions along the extrageniculate pathway, alongside reduced or unchanged responses within the geniculostriate pathway, compared to wild-type (WT) controls. To establish the pathological context of these functional changes, we quantified subunit c of mitochondrial ATP synthase (SCMAS), whose accumulation is a common pathological marker of CLN3 disease, across visual regions and found region-dependent differences in storage burden. Together, these findings demonstrate pathway-specific functional alterations in early CLN3 disease and suggest that enhanced extrageniculate activity may reflect a compensatory response to geniculostriate dysfunction. By identifying functional alterations at an early stage, this work provides insight into disease progression and highlights the potential of fUS as a promising tool for early detection of disease-related functional changes in Batten disease.

## Methods

### Animals

All procedures were performed in accordance with the guidelines of the National Institutes of Health (NIH) and were approved by the University Committee on Animal Resources (UCAR) at the University of Rochester.

*Cln3−/−* mice and age-matched wild-type (WT) littermates of both sexes were bred in-house from the Jackson Laboratory (B6.129S6-Cln3tm1Nbm/J /J, JAX:029471, ME). The mutant line was first generated on the 129S6 genetic background (Mitchison et al., 1999), and later backcrossed to the C57BL/6J background for 10 generations before deposition at the Jackson Laboratory. Genotypes were confirmed by Transnetyx (Cordova, TN) using a strain-specific real-time PCR assay according to the provider’s protocol.

Mice were housed under controlled temperature and humidity on a 12–12 h light-dark cycle, with unlimited access to food and water. fUS imaging was conducted in mice aged 12–14 weeks, including 10 WT animals (5 males and 5 females) and 12 *Cln3−/−* animals (5 males and 7 females). No signs of distress were observed during imaging sessions. Mice used in pathological analyses were all 3 months old and included 8 WT animals (4 males and 4 females) and 8 *Cln3−/−* animals (4 males and 4 females).

### Animal preparation

Anesthesia was induced using a gas induction chamber with 5% isoflurane mixed with medical oxygen. Animals were then positioned in a stereotaxic frame on a water-heating pad to maintain body temperature at approximately 37 °C. Anesthesia was maintained during surgical preparation with 1.5% isoflurane delivered via a nose cone, and ophthalmic ointment was applied to prevent corneal drying. Hair over the scalp was removed using a depilatory cream, followed by a midline scalp incision to expose a sufficient cranial area for imaging the visual pathway while leaving the skull intact. A 3% hydrogen peroxide (H_2_O_2_) solution was applied to remove residual membrane tissue from the skull surface. The exposed skull surface was then rinsed with sterile saline to ensure a clean acoustic interface. Following surgical preparation, the isoflurane concentration was reduced to 0.5%, and dexmedetomidine (0.03 mg/kg) was administered intraperitoneally (I.P.) to reduce muscle movement, using a protocol modified from an established fMRI anesthesia approach (Grandjean et al., 2014). Imaging began 30 minutes after dexmedetomidine administration. After fUS imaging, the scalp incision was sutured, and the antibiotic ointment was applied to the suture area. Animals were returned to individual cages once fully awake and freely moving and were closely monitored during recovery.

### Transcranial functional ultrasound imaging

Functional ultrasound imaging was performed using a programmable ultrafast ultrasound scanner (Vantage 128, Verasonics Inc., Kirkland, WA, USA) equipped with an L22–14vX linear array transducer (Verasonics Inc., Kirkland, WA, USA). The transducer operated at a center frequency of 15 MHz with a pulse repetition frequency (PRF) of 15 kHz. The transducer was mounted onto a stereotaxic frame using a custom-made probe holder. Probe positioning was controlled using a three-axis stereotaxic manipulator integrated with a digital readout to monitor probe displacement during imaging. Ultrasound gel was applied between the probe and the intact skull to ensure effective acoustic coupling, and the probe was positioned approximately 2 mm above the cortical surface in the coronal plane. After alignment with the coronal plane of the brain, the probe was moved exclusively along the anteroposterior axis. To cover the visual pathway, coronal planes spanning from Bregma −1.70 mm to Bregma −2.20 mm and from Bregma −2.80 mm to Bregma −3.70 mm were acquired. The optimal imaging planes were identified by visual inspection, matching real-time power Doppler images with a reference vascular atlas generated in-house. Consecutive coronal planes were acquired at 0.10 mm intervals. At each coronal plane, one visual stimulation session was presented while functional ultrasound images were acquired. The imaging depth was set from 0 to 8 mm, covering the full depth of the mouse brain. Ultrafast ultrasound acquisition was performed using compounded plane-wave transmissions (Montaldo et al., 2009) at 11 steering angles, uniformly spaced between −5° and +5°. Each power Doppler image was obtained from 200 compounded frames acquired at a frame rate of 1000 Hz. Channel data were acquired using a constant time-gain compensation (TGC) profile for further processing.

### Visual stimulation

Visual stimuli were presented on a 16-inch monitor positioned 30 cm from the right eye and oriented 45° relative to the anteroposterior axis of the mouse. The stimulus consisted of sinusoidal luminance gratings oriented at 45°, with an effective spatial period of approximately 0.13 cycles per degree based on the monitor geometry. Each session, corresponding to a single coronal slice, began with a 20-second black-screen baseline at zero luminance, followed by the grating simulation flashed at 2 Hz with a 0.5 duty cycle for 10 seconds, and a 15-second black-screen recovery period. A small, uniformly distributed temporal jitter (±50 *ms*) was added to the black phase to reduce strict periodicity and potential adaptation. This stimulation-recovery sequence was repeated 10 times per session. All visual stimulation paradigms used in this study were generated using custom-written MATLAB scripts (MathWorks, Natick, MA). Between sessions, the mice were kept in a dark environment, with an interval of at least 5 minutes between consecutive sessions.

### Ultrasound signal processing

B-mode image reconstruction was performed using a delay-and-sum beamforming algorithm to generate in-phase/quadrature (IQ) data from raw channel data. Delay-and-sum beamforming was implemented in MATLAB. Compounded B-mode images were obtained by coherently combining B-mode images from 11 steering angles. Singular value decomposition (SVD)-based spatiotemporal filtering (Demene et al., 2015) was then applied to the sequence of 200 compounded images to distinguish the tissue and blood signals. The leading singular value components corresponding to tissue signals were set to zero, and the remaining components were summed to calculate the pixel-wise intensity, forming the power Doppler image.

### Activation maps

Activation maps were computed to quantify fUS responses to visual stimulation using established approaches (Mace et al., 2011). After constructing power Doppler images, a temporal blood-flow signal can be extracted for each pixel. Time points identified as motion artifacts were excluded from further analysis.

To account for hemodynamic effects under light anesthesia, a predicted fUS response was generated by convolving the stimulus pattern with a standard hemodynamic response function (HRF) (Brunner et al., 2020; Brunner et al., 2021; Friston et al., 1998). The stimulus pattern was defined as a binary time series, with a value of 1 during the 10-s flashing-grating period and 0 during the black-screen period. The HRF was modeled as a combination of two gamma functions (Friston et al., 1998):

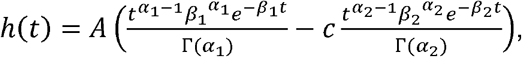

where *α*_l_ = 6, *α*_2_ = 16, *β*_l_ = *β*_2_ =1 and *c* = 1/6. Γ denotes the gamma function, and *A* is a scaling factor. These parameters correspond to commonly used HRF settings.

For each pixel, a Pearson correlation coefficient was computed between the pixel-wise temporal signal and the predicted fUS response, yielding a correlation map. Correlation values were converted to Z-score maps using Fisher’s transform, defined as

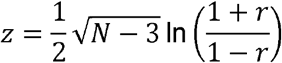

Only pixels exceeding a stringent Z-score threshold of 3.1 (corresponding to *p* < 0.001 in a one-tailed test) were considered activated to reduce false-positive detections (Woo et al., 2014). In addition, connected components smaller than 0.10 *mm* x 0.10 *mm* were removed based on the spatial resolution of fUS imaging. The activated regions were overlaid onto the corresponding grayscale power Doppler images to generate the activation maps shown in the figures.

### Registration

For each animal, consecutive coronal slices were stacked to reconstruct a three-dimensional blood-flow volume. This volume was manually registered to the Allen Mouse Brain Common Coordinate Framework (CCFv3) (Wang et al., 2020) using the open-source software 3D Slicer (Fedorov et al., 2012). The CCFv3 template and annotation volumes were obtained from the Scalable Brain Atlas (Bakker et al., 2015). Three-dimensional affine transformations, including scaling, translation, and rotation, were applied to the blood-flow volume to align anatomical landmarks and boundaries with the atlas. Following registration, the blood-flow volume and activation maps were projected onto the corresponding atlas-defined brain regions. Registration accuracy was verified by visual inspection of anatomical correspondence between the atlas and the blood-flow volume. Using the registered volumes, activation information were extracted from the following regions of interest (ROI): lateral geniculate complex (LGN), lateral posterior nucleus of the thalamus (LP), superior colliculus (SC), primary visual area (V1), anterolateral visual area (AL), posteromedial visual area (PM), lateral visual area (L) and pretectal nuclei (PN), including nucleus of the optic tract (NOT), posterior pretectal nucleus (PPT) and anterior pretectal nucleus (APN). The prefixes “a” and “p” were used to denote the anterior and posterior subdivisions of the ROI, respectively.

For anatomical organization, some ROIs were grouped for analysis. The anterior thalamus (aTH) comprised the LGN and aLP, while the posterior thalamus (pTH) corresponded to the pLP. The anterior midbrain (aMB) included aSC, NOT, PPT, and APN, and the posterior midbrain (pMB) referred to pSC. Visual cortical areas were analyzed as separate ROIs. For pathway-level organization and visualization, secondary visual cortex (V2) was referenced collectively, with anterior V2 (aV2) denoting aPM and AL, and posterior V2 (pV2) denoting pPM and L.

### fUS statistical analysis

To quantify activation strength from fUS data, two complementary metrics, extent and intensity, were computed for each region of interest (ROI) using custom-written MATLAB scripts. Data were pooled across coronal slices within predefined anteroposterior coordinate ranges. Activation extent was quantified as the percentage of activated pixels, defined as the ratio of pixels exceeding the activation threshold within an ROI to the total number of pixels in that ROI. Because this metric is bounded and non-normally distributed, the square-root transform was applied to the percentage of activated pixels for visualization, reducing compression near zero and improving dynamic range. This transformation was applied solely for display purposes and did not affect statistical inference, which was conducted using non-parametric methods. Activation intensity was quantified as the mean Z-score within each ROI, obtained by averaging Z-scores across all pixels within the defined region, providing a measure of response magnitude.

Statistical analyses were performed using GraphPad Prism 10.6.1 (GraphPad Software, Boston, MA, USA) and MATLAB, with data visualization generated in GraphPad Prism. For statistical analysis of the fUS data, each mouse contributed one value per defined ROI (including grouped regions where applicable) for each activation metric, computed within the selected coronal imaging window. Genotype-dependent differences were first assessed at the ROI-level. Normality was evaluated using the Shapiro–Wilk test. For the percentage of activated pixels, data in most ROIs failed normality tests. Therefore, two-sided unpaired Mann-Whitney tests were performed for each ROI, and results are presented as medians with interquartile ranges (IQRs). Mean Z-score values met normality assumptions in each ROI, and therefore were analyzed using two-sided unpaired t-tests, with results reported as mean ± SEM.

To evaluate genotype effects at the pathway level, ROIs were assigned to visual pathways based on known anatomical connectivity. The geniculostriate pathway included the anterior thalamus (aTH), V1, and pV2, whereas the extrageniculate pathway included the midbrain (MB), posterior thalamus (pTH), and aV2. Mixed-effect analyses were performed separately for each activation metric and pathway. For activation extent, a binomial generalized linear mixed-effect model with a logit link function was used in MATLAB. For activation intensity, mixed-effects analysis was performed in GraphPad Prism. In these models, genotype, ROI, and ROI × Genotype interaction were treated as fixed effects, with subject included as a random effect. When multiple comparisons were performed within the mixed-effects framework, P values were adjusted using Tukey’s post hoc test to control for family-wise error.

Statistical significance was defined as *p* < 0.05, and values between 0.05 and 0.10 were considered to indicate a statistical trend.

### Histology and Immunohistochemistry

#### Perfusion and sectioning

Animals were anesthetized with ketamine/xylazine (K/X; K: 100 mg/kg; X: 10 mg/kg) and perfused with 1-times concentration of phosphate-buffered saline (1X PBS) and 4% paraformaldehyde (PFA). Brains were dissected and fixed in 4% PFA for 24 hours before being stored in 1X PBS with 0.1% sodium azide (NaN_3_) at 4°C. Brain tissue was sectioned coronally at a thickness of 80μm using a VF-310-0Z Compresstome (Precisionary Instruments, MA). Slices were stored in 1X PBS with 0.1% NaN_3_ at 4°C in 24-well tissue culture plates (Thermo Fisher Scientific Inc., MA). Brain sections from both *Cln3−/−* and WT mice were collected at 3 months of age.

#### Immunostaining

Brain sections containing the thalamus, midbrain, and visual cortex were selected for immunostaining to assess central visual neuropathology. Immunostaining was performed on free-floating brain sections in 24-well plates using two to three slices per well. Slices were first washed in 1X PBS three times for 5 minutes. Non-specific protein binding was blocked using 5% normal goat serum (NGS) in 1X PBS with 0.01% Triton X-100 (PBS-T) for 1 hour. After blocking, slices were incubated overnight at 4°C in an antibody against SCMAS (1:250, anti-rabbit, ab181243, Abcam, MA) diluted in 5% NGS + PBS-T. To ensure robust visualization of brain cells, slices were washed 3 x 5 minutes in 1X PBS and incubated for 5 minutes in DAPI (nuclei) (1:100 from 300µM intermediate solution, D1306, Thermo Fisher Scientific Inc., MA) diluted in 1X PBS. Lastly, slices were washed 3 x 5 minutes in 1X PBS, mounted with media without DAPI (InvitrogenTM ProLongTM Glass Antifade Mountant, P36982, Thermo Fisher Scientific Inc., MA), and coverslipped on slides.

#### Imaging

After immunostaining, brain sections were imaged by an Olympus VS120 Slide Scanner (Evident Scientific, PA) at the Center for Musculoskeletal Research (CMSR) (URMC, NY) and a Nikon A1R HD – Pikachu Confocal Microscope (Nikon Corporation, Japan) at the Center for Advanced Microscopy & Nanoscopy (CALMN) (URMC, NY). The slide scanner was used to quickly compare storage material accumulation at low magnification and resolution. Confocal microscopy was used to assess SCMAS expression in specific areas at higher magnification and resolution. Confocal images were acquired at 20X (0.4 µm per pixel) for the thalamus, midbrain, and visual cortex at blue (DAPI) and red (SCMAS) channels.

#### Imaging analysis

Imaging analysis was performed to quantify SCMAS expression using customized MATLAB scripts. For SCMAS quantification, images from the SCMAS channel were background-subtracted, thresholded using Sauvola’s adaptive thresholding algorithm, and subsequently binarized. Standard global thresholding methods, such as Otsu’s algorithm, were not applied because SCMAS signals in WT mice were sparse and low-intensity, violating the bimodal distribution assumption. SCMAS expression was quantified as SCMAS^+^ area per cell (SCMAS^+^ area per cell = total SCMAS^+^ area above the threshold / total number of cells; cell number determined by DAPI).

#### Statistics

Statistical analyses and data visualization were performed using GraphPad Prism 10.6.1. Each mouse contributed one value per ROI. The ROIs were categorized into three groups: thalamus, visual cortex, and midbrain. For each group, a two-way ANOVA was conducted to evaluate the effects of genotypes and ROIs, followed by multiple comparisons. When more than two ROIs were in the group, Tukey’s correction was applied to the multiple comparisons. Data were presented as mean ± SEM. Significance was defined as *p* < 0.05.

## Results

### WT mice showed stimulus-evoked fUS activation in visual areas

To assess the reliability of functional ultrasound (fUS) for detecting stimulus-evoked activity and establish baseline visual responses, activation patterns were examined in young adult (3-month-old) wild-type (WT) mice across coronal planes spanning major components of the visual pathway (Fig. 1). At Bregma −2.00 mm, clear and spatially localized activation was observed in the lateral posterior nucleus (LP) and the lateral geniculate nucleus (LGN). At Bregma −3.00 mm, activation patterns were more spatially heterogeneous, with detectable responses in the primary visual cortex (V1) and the superior colliculus (SC), accompanied by weaker and more diffuse activation in the anterolateral visual area (AL) and the posteromedial visual area (PM). At Bregma −3.40 mm, pronounced and spatially coherent activation was observed in both V1 and SC. Across these coronal locations, activation patterns were reproducible across animals and anatomically aligned with known visual pathway structures, demonstrating the sensitivity of fUS for mapping stimulus-evoked visual responses. These WT activation maps therefore provide a stable reference for subsequent comparisons between WT and *Cln3−/−* mice.

**Figure 1.**
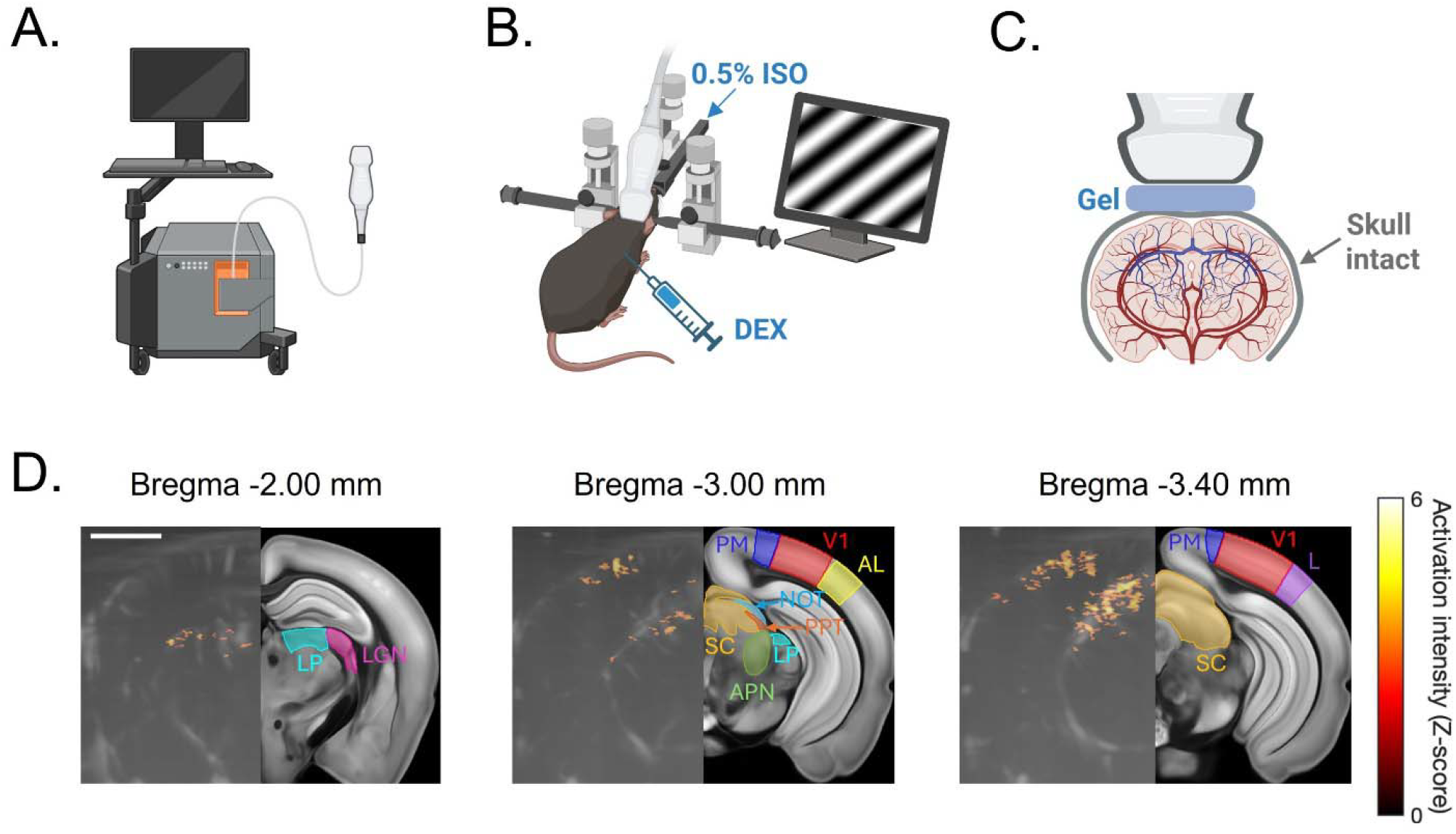
Experimental setup and representative functional ultrasound (fUS) responses in a wild-type mouse. (A) Schematic of the fUS imaging system. (B) Experimental configuration showing a mouse positioned in a stereotaxic frame during imaging. The ultrasound probe was placed on top of the head, and a monitor for visual stimulus presentation was positioned on the right side of the animal. Anesthesia (0.5% isoflurane) was adjusted prior to imaging, and a bolus injection of dexmedetomidine (DEX) was administered to minimize movement. (C) Coronal schematic illustrating the intact-skull imaging configuration. Ultrasound gel was applied between the skull and the probe for acoustic coupling. (D) Representative fUS activation maps from a wild-type (WT) mouse at three coronal planes corresponding to regions included in the quantitative analyses. In each panel, activated pixels (color-coded by Z-score) are overlaid on the grayscale blood flow image (left), with the corresponding anatomical reference and region labels shown on the right. At Bregma −2.00 mm, activation is observed in the lateral posterior nucleus (LP) and lateral geniculate nucleus (LGN). At Bregma −3.00 mm, activation is detected in primary visual cortex (V1) and superior colliculus (SC), with weaker responses in secondary visual cortex. At Bregma −3.40 mm, activation is evident in V1 and SC along the primary visual pathway. Scale bar: 2 mm.

### *Cln3−/−* mice showed altered fUS activation across various visual areas

To examine the effects of the *Cln3* mutation on visual processing, fUS activation was used to compare young adult (3-month-old) *Cln3−/−* mice and WT controls across major nodes of the visual system, including thalamic nuclei, cortical areas, and midbrain structures. Activation maps provided a qualitative overview of response patterns, while ROI-based statistical analyses further quantitatively confirmed genotype-dependent differences.

Within the visual thalamus, opposing alterations in activation were observed between anterior and posterior subdivisions (Fig. 2A–B). Specifically, *Cln3*−/− mice exhibited increased activation in the posterior thalamus (pTH) and reduced activation in the anterior thalamus (aTH). ROI-level Mann–Whitney tests revealed a significantly higher percentage of activated pixels in the pTH of *Cln3−/−* mice compared to WT animals (Fig. 2C; *p* = 0.0370). In addition, *Cln3−/−* mice showed a significantly attenuated response magnitude in aTH, reflected by a reduced mean Z-score (Fig. 2D; *p* = 0.0315, unpaired t-test).

**Figure 2.**
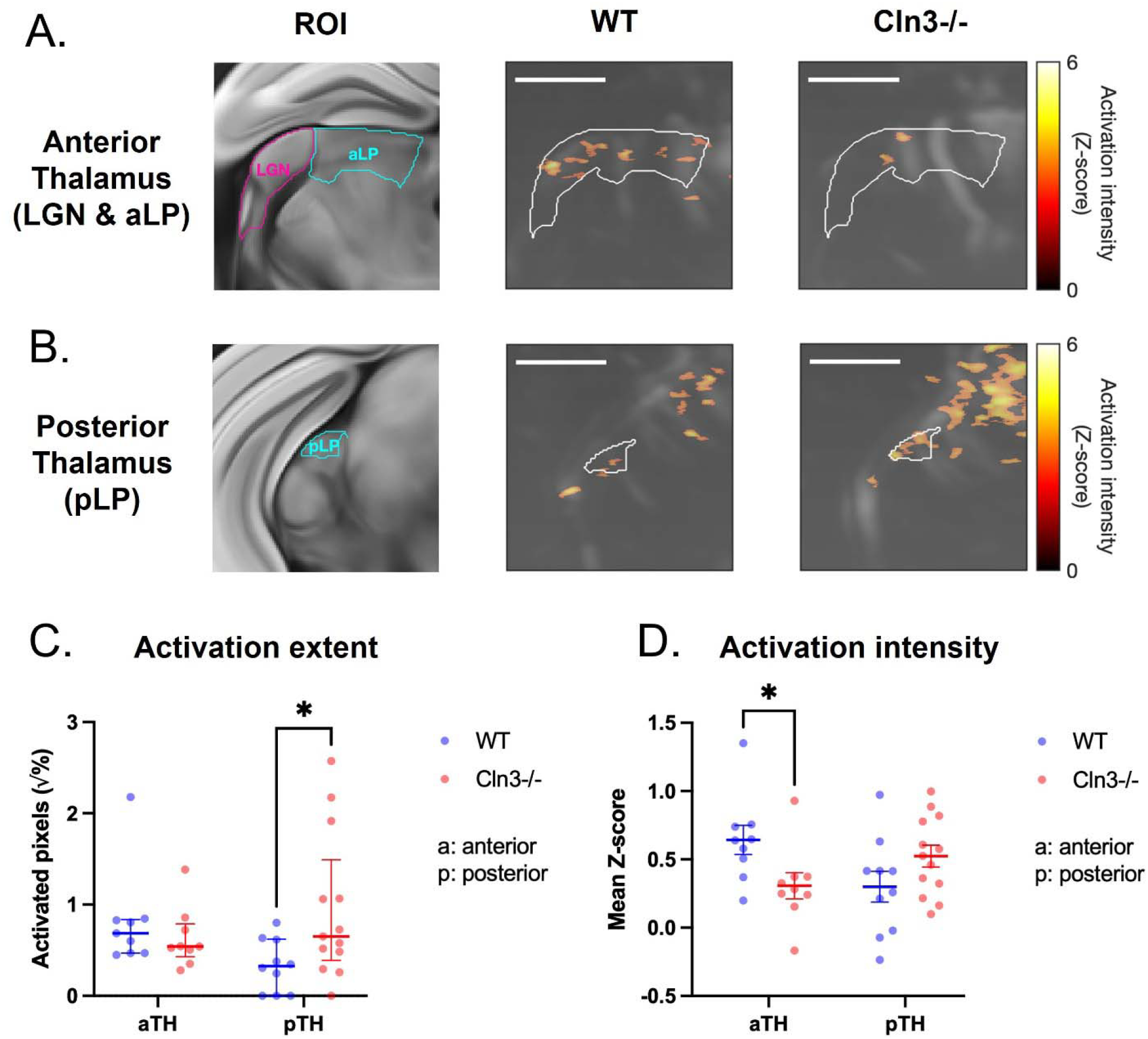
Activation differences in the visual thalamus between WT and *Cln3−/−* mice. (A–B) Coronal anatomical reference images of visual thalamic regions of interest (ROIs), including LP and LGN, with representative fUS activation maps from WT (left) and *Cln3−/−* (right) mice. (A) Anterior visual thalamus (aTH), including LGN and aLP, at Bregma −2.00 mm. (B) Posterior visual thalamus (pTH) at Bregma −3.00 mm. Scale bars: 1 mm. (C) Square root–transformed percentage of activated pixels within each ROI. Each dot represents one animal (WT, blue; *Cln3−/−*, red). Sample sizes were WT, *n* = 10 and *Cln3−/−, n* = 13, except for aTH (WT, *n* = 9; *Cln3−/−, n* = 9). Bars the Mann–Whitney U test. A significant difference was observed in pTH (*p* = 0.0379). indicate median and interquartile range. Genotype comparisons were performed using animal (WT, blue; *Cln3−/−*, red). Sample sizes were WT, *n* = 10 and *Cln3−/−, n* = 13, (D) Mean Z-score of activation intensity within each ROI. Each dot represents one except for aTH (WT, *n* = 9; *Cln3−/−, n* = 9). Bars indicate mean ± SEM. Genotype comparisons were performed using a two-sided unpaired t-test. A significant difference was observed in aTH (*p* = 0.0315). Non-significant *p*-values are not shown.

Across visual cortical areas, altered activation patterns were most evident in anterior regions (Fig. 3). ROI-based comparisons demonstrated significantly higher percentages of activated pixels in aPM and AL of *Cln3*−/− mice relative to WT controls (Fig. 3C; aPM: *p* = 0.0254, AL: *p* = 0.0303, Mann-Whitney test), consistent with the more extensive activated clusters observed in anterior cortical activation maps (Fig. 3A– B). Posterior cortical regions did not show significant genotype-dependent changes in either the percentage of activated pixels or the mean Z-score (Fig. 3C–D).

**Figure 3.**
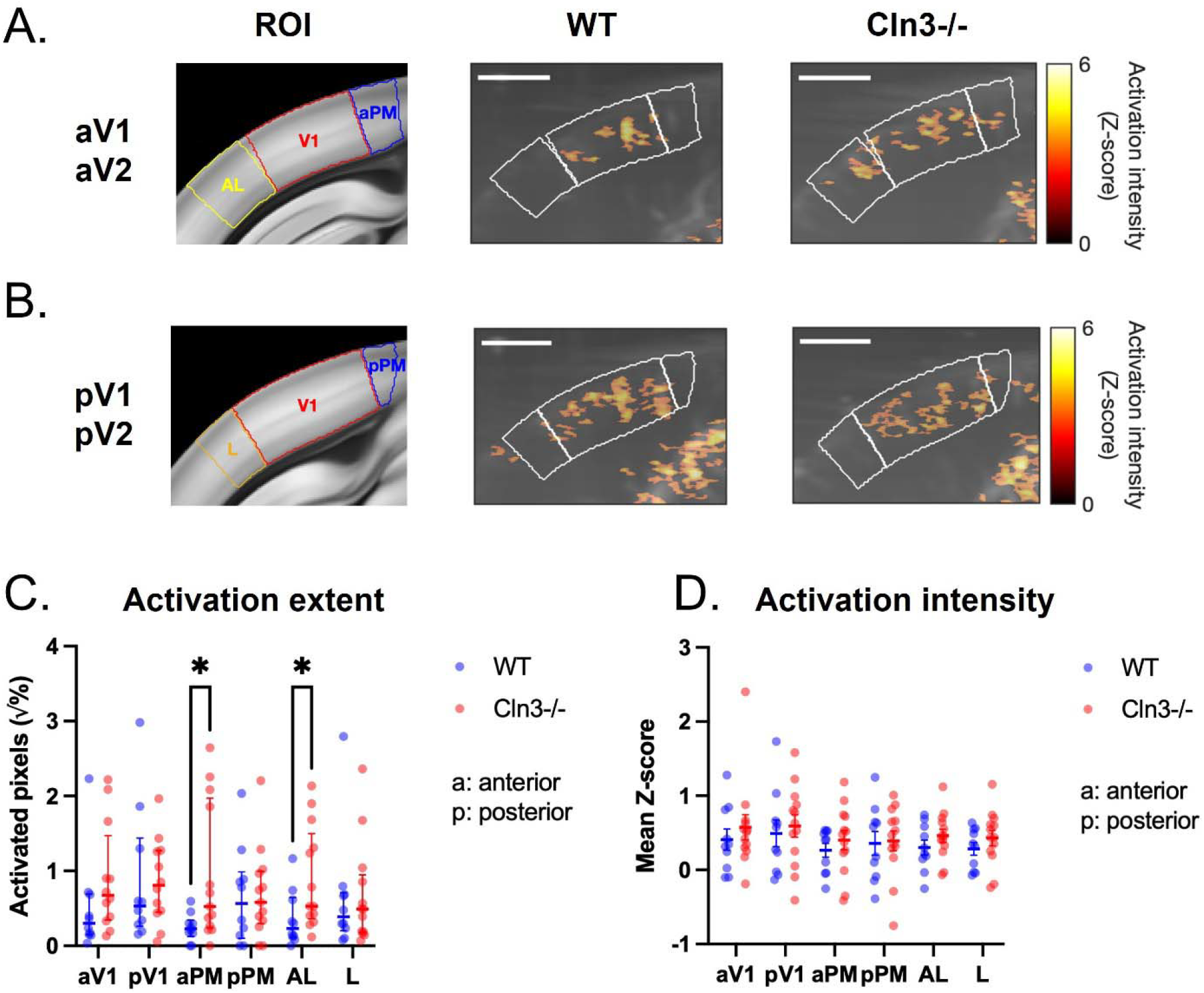
Activation differences in visual cortices between WT and *Cln3−/−* mice. (A–B) Coronal anatomical reference images of primary and higher visual cortical ROIs, including V1, PM, AL, and lateral visual cortex (L), with representative fUS activation maps from WT (left) and *Cln3−/−* (right) mice. (A) Anterior visual cortex, at Bregma −3.00 mm. (B) Posterior visual cortex at Bregma −3.40 mm. Some ROIs were subdivided into anterior (a) and posterior (p) regions for quantitative analysis. Scale bars: 1 mm. (C) Square root–transformed percentage of activated pixels within each ROI. Each dot represents one animal (WT, blue, *n* = 10; *Cln3−/−*, red, *n* = 13). Bars indicate median Whitney U test. Significant differences were observed in aPM (*p* = 0.0254) and aAL and interquartile range. Genotype comparisons were performed using the Mann– (*p* = 0.0303). (D) Mean Z-score within each ROI. Each dot represents one animal (WT, blue, *n* = 10; *Cln3−/−*, red, *n* = 13). Bars indicate mean ± SEM. Genotype comparisons were performed using a two-sided unpaired t-test. Non-significant *p*-values are not shown.

In the midbrain, genotype-related differences were most pronounced in the anterior regions (Fig. 4). Direct genotype comparisons revealed a significantly higher percentage of activated pixels in the anterior midbrain (aMB) of *Cln3*−/− mice compared to WT animals (Fig. 4C; *p* = 0.0358, Mann-Whitney test), corresponding to the expanded activation observed in the anterior midbrain (Fig. 4A). No significant genotype-dependent differences were detected in the posterior midbrain or in activation intensity in either subdivision.

**Figure 4.**
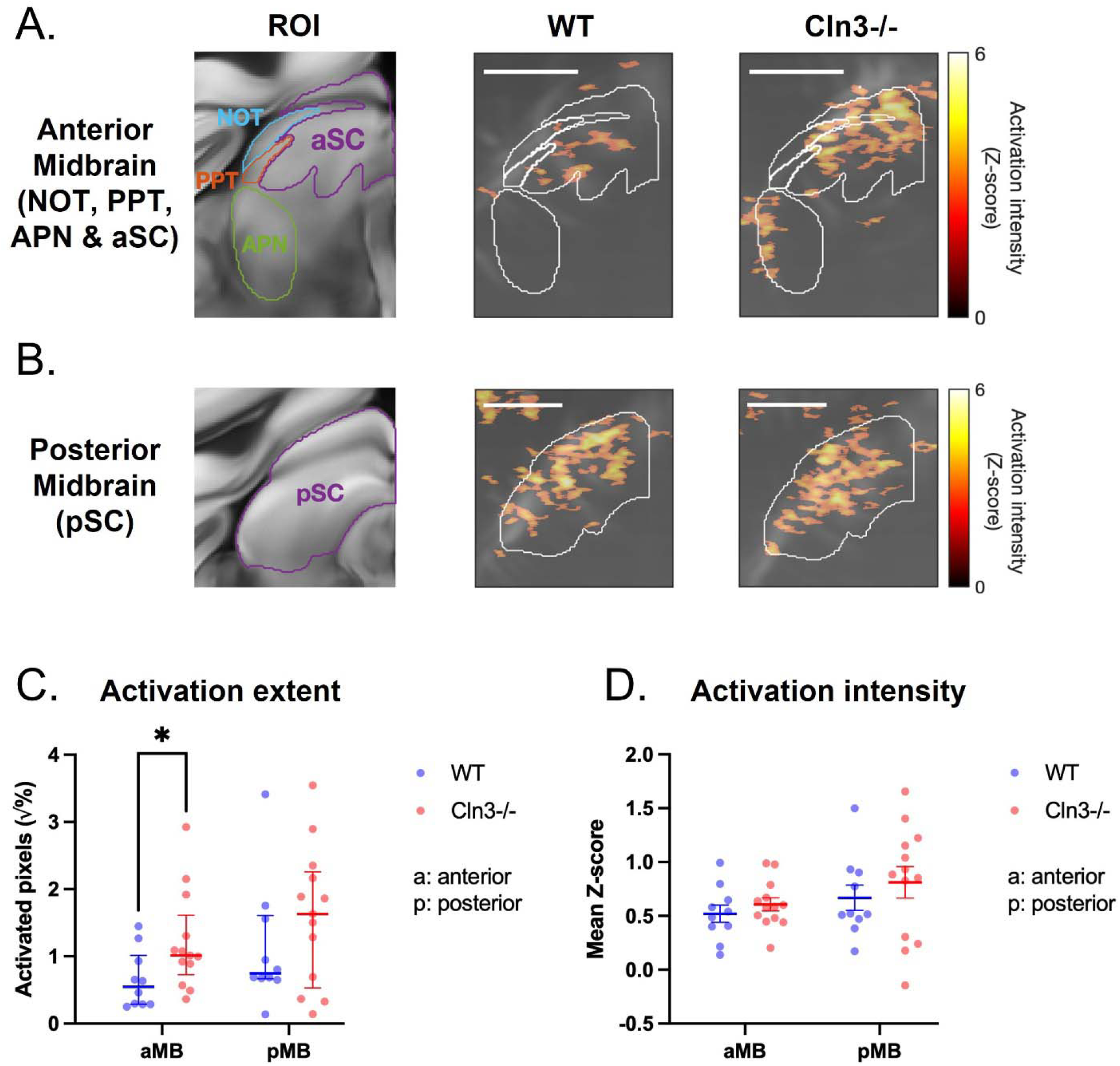
Activation differences in the midbrain between WT and *Cln3−/−* mice. (A– B) Coronal anatomical reference images of anterior and posterior midbrain ROIs, including the SC, NOT, PPT, and APN, with representative fUS activation maps from WT (left) and *Cln3−/−* (right) mice. (A) Anterior midbrain (aMB) at Bregma −3.00 mm. (B) Posterior midbrain (pMB) at Bregma −3.40 mm. Scale bars: 1 mm. (C) Square root– transformed percentage of activated pixels within each ROI. Each dot represents one animal (WT, blue, *n* = 10; *Cln3−/−*, red, *n* = 13). Bars indicate median and interquartile significant difference was observed in aMB (*p* = 0.0358). (D) Mean Z-score within range. Genotype comparisons were performed using the Mann–Whitney U test. A each ROI. Each dot represents one animal (WT, blue, *n* = 10; *Cln3−/−*, red, *n* = 13). Bars indicate mean ± SEM. Genotype comparisons were performed using a two-sided unpaired t-test. Non-significant *p*-values are not shown.

### Region-specific differences in SCMAS accumulation in *Cln3−/−* mice

To provide a pathological framework for interpreting the functional alterations detected by fUS, SCMAS expression was quantified across visual thalamic, cortical, and midbrain regions. Across all regions of interest, *Cln3−/−* mice exhibited significantly higher SCMAS+ area per cell than WT animals, indicating a pervasive pathological burden throughout the visual system. Notably, the extent of SCMAS accumulation varied substantially across regions within *Cln3−/−* mice. Therefore, rather than reiterating genotype comparisons for each ROI, subsequent analyses focused on characterizing regional differences in SCMAS accumulation within the *Cln3−/−* group, providing a structural context for interpreting the heterogeneous functional changes detected by fUS.

Within the visual thalamus, SCMAS accumulation differed significantly between anterior and posterior subdivisions in *Cln3−/−* mice (Fig. 5). Two-way ANOVA revealed significant main effects of genotype (*F* (1,28) = 10.8, *p* = 0.0028) and ROI (*F* (1,28) = 1624, *p* < 0.0001), as well as a significant genotype × ROI interaction (*F* (1,28) = 13.8, *p* = 0.0009). Multiple comparisons showed that, within *Cln3−/−* mice, aTH exhibited significantly greater SCMAS+ area per cell than pTH (*p* < 0.0001), whereas no anterior-posterior difference was observed in WT animals (*p* = 0.7590). These findings suggest more severe pathology in the anterior thalamus than the posterior thalamus in *Cln3−/−* mice.

**Figure 5.**
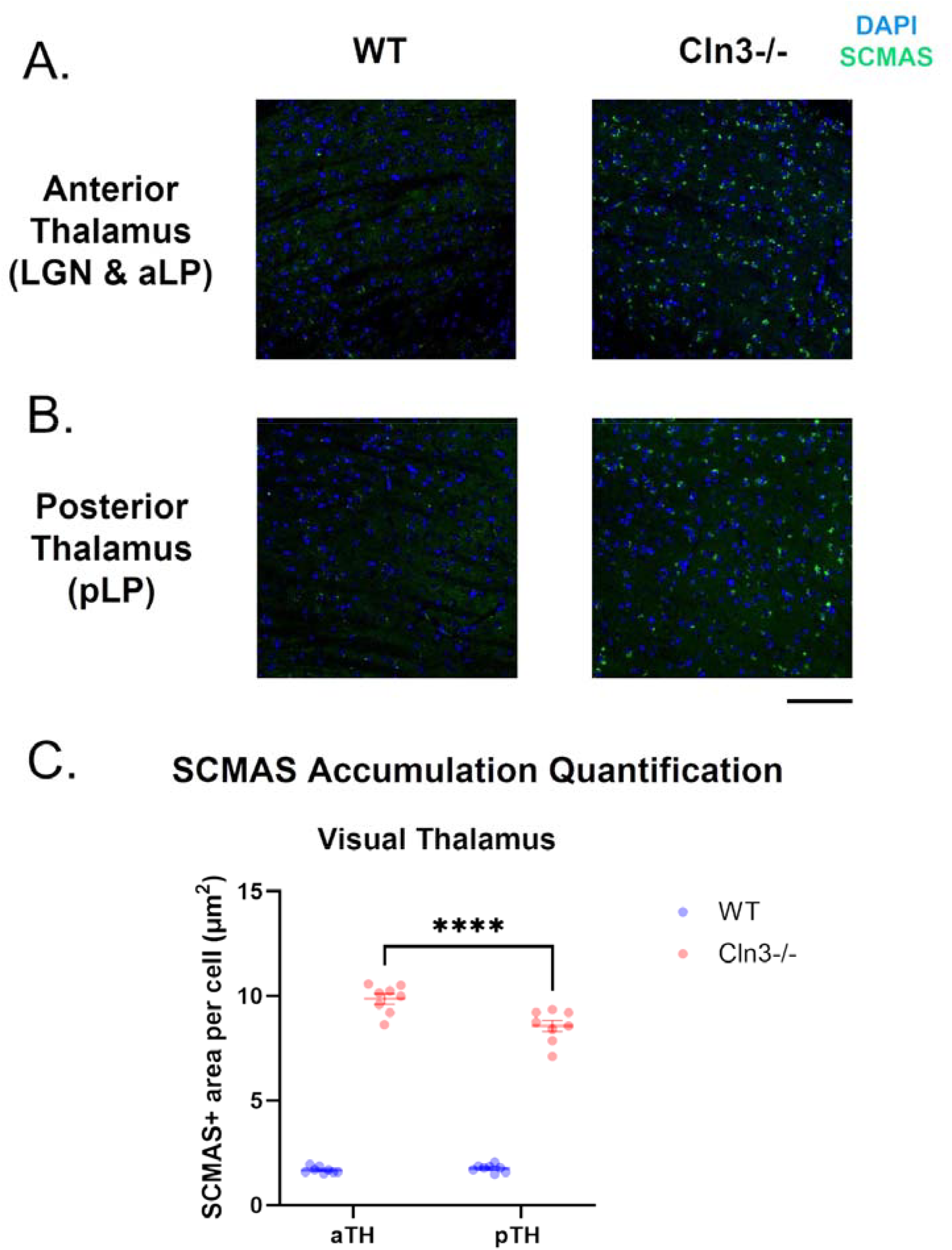
SCMAS accumulation in the visual thalamus of WT and *Cln3−/−* mice. (A-B) Representative immunofluorescence images of anterior and posterior visual thalamic regions from WT (left) and *Cln3−/−* (right) mice. Blue dots represent DAPI-labeled nuclei, and green punctate clusters indicate SCMAS accumulation. Scale bar: 100 μm. (C) animal (WT, blue, *n* = 8; *Cln3−/−*, red, *n* = 8), and bars indicate mean ± SEM. Quantification of SCMAS+ area per cell (μm^2^) in aTH and pTH. Each dot represents one Statistical analysis was performed using two-way ANOVA followed by multiple regions (*p* < 0.0001). Within *Cln3−/−* mice, SCMAS accumulation was significantly higher comparisons. SCMAS accumulation was higher in *Cln3−/−* mice than in WT mice for both in aTH than pTH (*p* < 0.0001).

In the visual cortex, regional differences in SCMAS were examined across primary and secondary visual areas (Fig. 6). Two-way ANOVA revealed significant main effects of ROI (*F* (3,56) = 7.1, *p* = 0.0004), genotype (*F* (1,56) = 2062, *p* < 0.0001), and a significant genotype × ROI interaction (*F* (3,56) = 7.6, *p* = 0.0002). Tukey-corrected multiple comparisons demonstrated that in *Cln3−/−* mice, both aV1 and pV1 showed significantly higher SCMAS+ area per cell than aV2 and pV2 (aV1 vs. aV2: *p* = 0.0005, aV1 vs. pV2: *p* = 0.0001, pV1 vs. aV2: *p* = 0.0003, pV1 vs. pV2: *p* < 0.0001), but no significant anterior-posterior differences were observed within V1 and V2. No regional differences were detected among cortical regions in WT animals. These results demonstrated that pathological burden is more elevated in V1 relative to V2.

**Figure 6.**
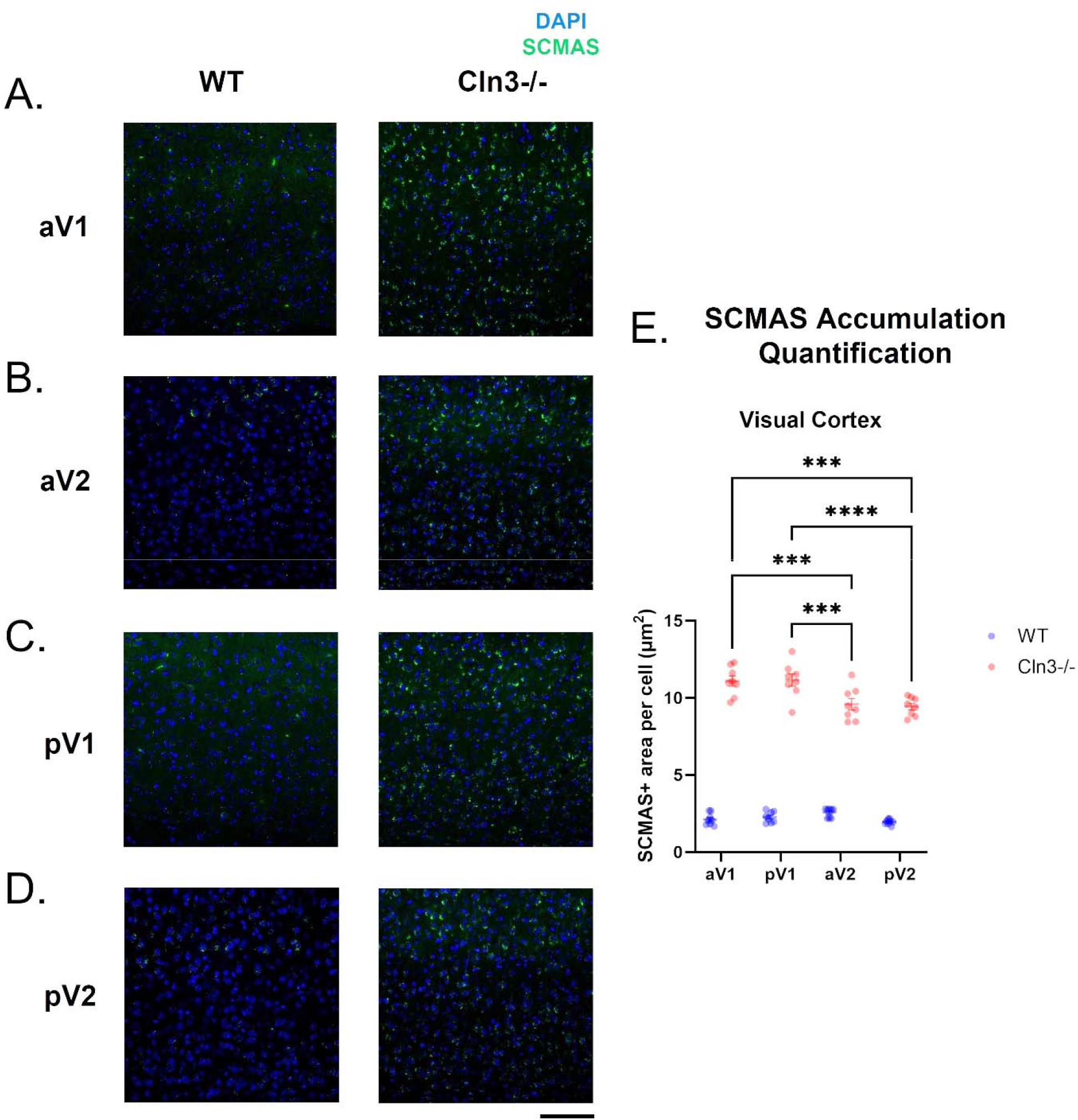
SCMAS accumulation in the visual cortex of WT and *Cln3−/−* mice. (A-D) Representative immunofluorescence images of anterior and posterior regions of V1 and V2 from WT (left) and *Cln3−/−* (right) mice. Blue dots represent DAPI-labeled nuclei, and green punctate clusters indicate SCMAS accumulation. Scale bar: 100 μm. (E) animal (WT, blue, *n* = 8; *Cln3−/−*, red, *n* = 8), and bars indicate mean ± SEM. Quantification of SCMAS+ area per cell (μm^2^) in each region. Each dot represents one Statistical analysis was performed using two-way ANOVA followed by Tukey’s multiple all regions (*p* < 0.0001). Within *Cln3−/−* mice, aV1 showed significantly higher SCMAS comparisons test. SCMAS accumulation was higher in *Cln3−/−* mice than in WT across levels than aV2 and pV2 (aV1 vs. aV2: *p* = 0.0005; aV1 vs. pV2: *p* < 0.0001), and pV1 showed higher levels than aV2 and pV2 (pV1 vs. aV2: *p* = 0.0003; pV1 vs. pV2: *p* < 0.0001). No significant anterior–posterior differences were observed within V1 or V2.

At the level of the midbrain, SCMAS accumulation differed between anterior and posterior subdivisions (Fig. 7). Two-way ANOVA revealed significant main effects of ROI (*F* (1,28) = 13.53, *p* = 0.0010), genotype (*F* (1,28) = 215.9, *p* < 0.0001), and a significant genotype × ROI interaction (*F* (1,28) = 7.57, *p* = 0.0103). In *Cln3−/−* mice, aMB, including aSC and pretectal nuclei, showed significantly higher SCAMS+ area per cell than pMB (*p* < 0.0001), whereas no regional difference was detected in WT animals. Notably, overall SCMAS levels in both midbrain regions were substantially lower than those observed in visual thalamic and cortical areas, indicating that the midbrain is comparatively less affected by pathological burden.

**Figure 7.**
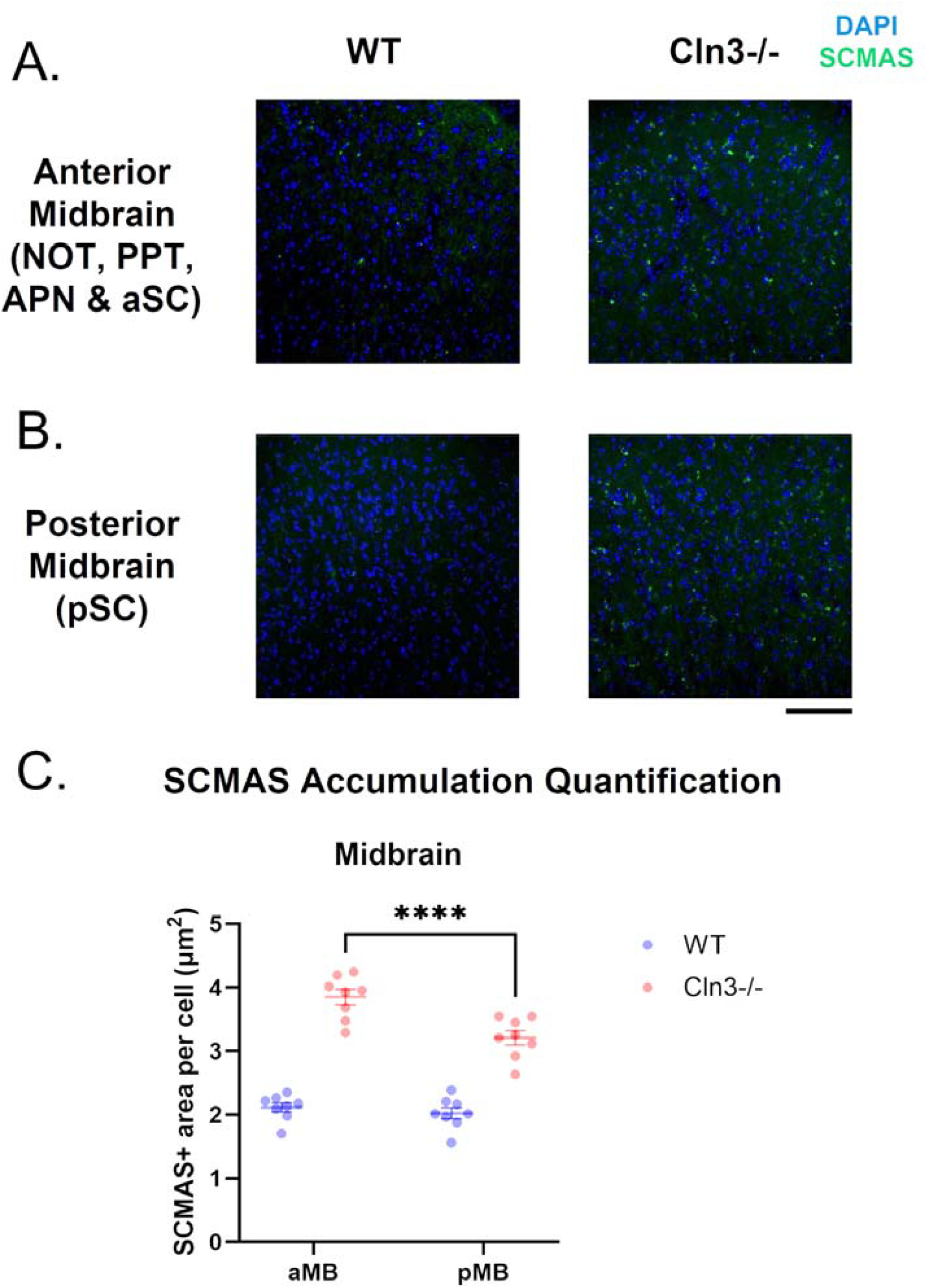
SCMAS accumulation in the midbrain of WT and *Cln3−/−* mice. (A-B) Representative immunofluorescence images of anterior and posterior midbrain from WT (left) and *Cln3−/−* (right) mice. Blue dots represent DAPI-labeled nuclei, and green punctate clusters indicate SCMAS accumulation. Scale bar: 100 μm. (C) Quantification blue, *n* = 8; *Cln3−/−*, red, *n* = 8), and bars indicate mean ± SEM. Statistical analysis of SCMAS+ area per cell (μm^2^) in each region. Each dot represents one animal (WT, accumulation was higher in *Cln3−/−* mice than in WT in both regions (*p* < 0.0001). Within was performed using two-way ANOVA followed by multiple comparisons. SCMAS *Cln3−/−* mice, SCMAS accumulation was significantly greater in aMB than in pMB (*p* < 0.0001).

### Pathway-specific relationships between fUS activation and SCMAS accumulation

To integrate the fUS and pathological findings across the visual system, pathway-level alterations were summarized in Fig. 8A. Within the geniculostriate pathway, *Cln3−/−* mice exhibited reduced activation in aTH relative to WT animals, with no significant differences observed in V1 or pV2. In contrast, *Cln3−/−* mice demonstrated significantly increased activation along the extrageniculate pathway, including midbrain, pTH, and aV2. Notably, regions in the extrageniculate pathway exhibited relatively lower SCMAS accumulation, with the midbrain showing the lowest pathological burden among the examined regions.

**Figure 8.**
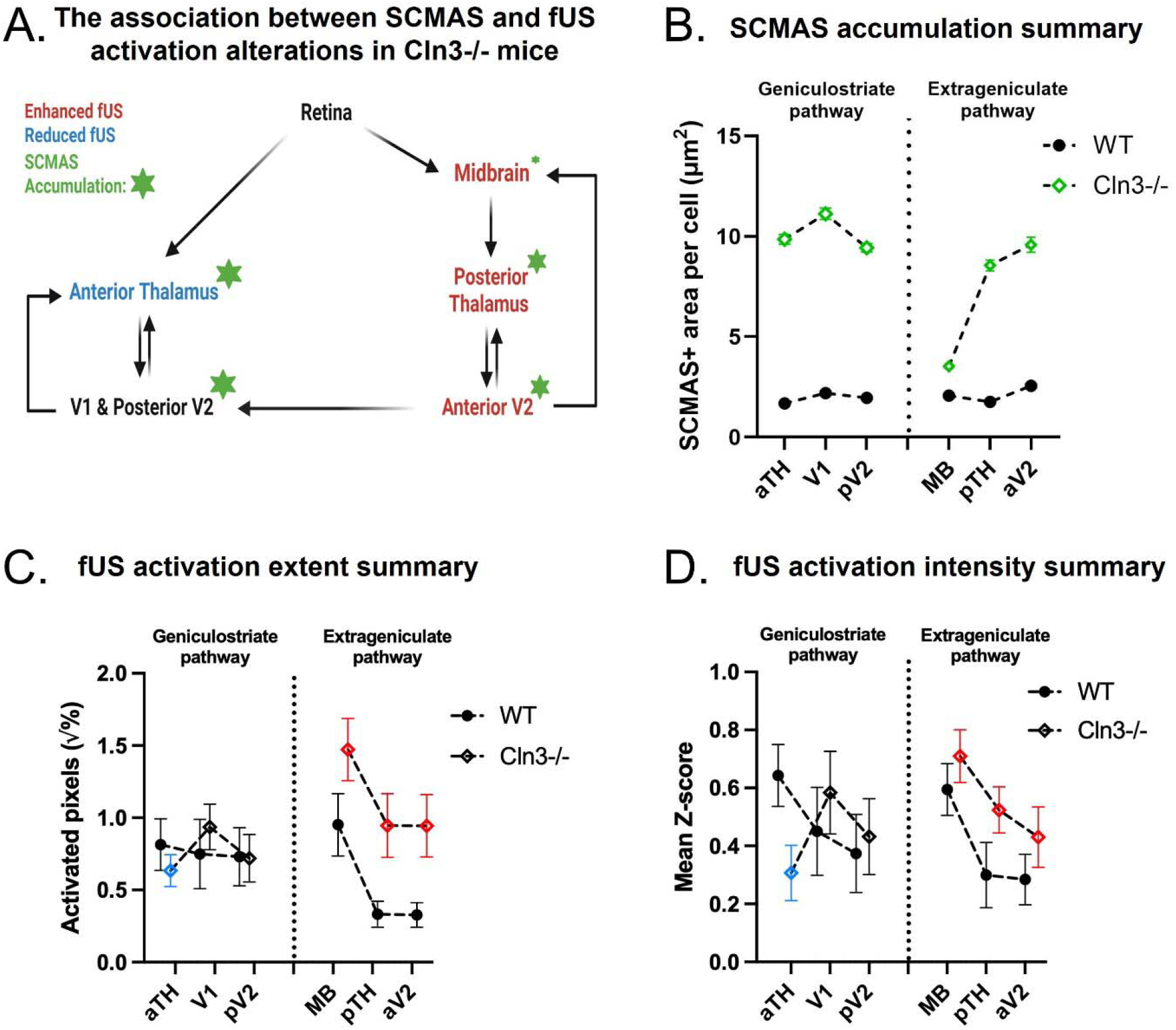
Pathway-level relationship between SCMAS accumulation and visually evoked activation in *Cln3−/−* mice. (A) Schematic summary of functional and pathological alterations along the two visual pathways. Arrows indicate major anatomical projections between regions. Red and blue labels denote regions showing increased or reduced activation in *Cln3−/−* mice relative to WT. Sizes of green stars represent relative levels of SCMAS accumulation. (B) SCMAS accumulation across visual regions in WT and *Cln3−/−* mice. WT and *Cln3−/−* are represented by filled circles and open diamonds, respectively. Marker size for *Cln3−/−* indicates relative SCMAS accumulation levels. The dotted line separates the two visual pathways. Values represent the SCMAS+ area per cell. Bars indicate mean ± SEM. (C-D) Visually evoked fUS activation metrics across visual regions in WT and *Cln3−/−* mice. WT and *Cln3−/−* are represented by filled circles and open diamonds, respectively. The dotted line separates the two visual pathways. Red and blue markers indicate regions with increased or decreased fUS response in *Cln3−/−* mice relative to WT. (C) Square root–transformed percentage of activated pixels. (D) Mean Z-score. Bars indicate mean ± SEM.

To facilitate pathway-level comparisons of pathological burden and functional activation between genotypes, SCMAS accumulation and visually evoked fUS responses from both WT and *Cln3−/−* groups are shown in Fig. 8B-D, with regions arranged according to visual pathway organization. SCMAS pathology was markedly elevated in *Cln3−/−* mice across all examined regions relative to WT animals (Fig. 8B). The greatest accumulation was observed in V1, which exceeded pV2, aTH, and pTH (*p* = 0.0293,*p* = 0.0483, and *p* = 0.0017, respectively). aTH also showed greater accumulation than pTH (*p* = 0.0008), whereas MB exhibited the lowest accumulation among all examined regions (all *p* < 0.0001). This regional pattern was supported by one-way ANOVA (*F* (3.155,22.08) = 101.2, *p* < 0.0001) with Tukey’s multiple comparisons.

fUS activation metrics also revealed pathway-specific functional alterations (Fig. 8C-D). To statistically assess pathway-level differences, mixed-effects analyses were performed separately for the two visual pathways (Supplementary Table 1). Within the extrageniculate pathway, generalized linear mixed-effects modeling of activation extent revealed significant main effects of ROI (*F* (4,105) = 5775.5, *p* < 0.0001), genotype (*F* (1,105) = 4.8610, *p* = 0.0030), and ROI × Genotype interaction (*F* (4,105) = 1796.7, *p* < 0.0001). Tukey’s multiple comparisons demonstrated increased activation extent in *Cln3−/−* mice in pTH, aPM, and AL (adjusted *p* = 0.0497,0.0356,and 0.0460, respectively), with a similar trend observed in aMB (adjusted *p* = 0.0727). Average activation intensity within the extrageniculate pathway similarly showed an increase in *Cln3−/−* mice compared to WT mice, although this effect did not reach statistical significance. In contrast, no significant genotype effects on activation extent were detected within the geniculostriate pathway. However, Tukey’s multiple comparisons of mean Z-score activation intensity within the geniculostriate pathway demonstrated significantly reduced activation in aTH in *Cln3−/−* mice (*p* = 0.0321).

Collectively, these analyses reveal a pathway-organized pattern of dysfunction in *Cln3−/−* mice. Regions along the extrageniculate pathway generally exhibited lower SCMAS accumulation together with enhanced visually evoked fUS activation, whereas regions within the geniculostriate pathway showed greater pathological burden and activation responses that were either unchanged or reduced relative to WT animals.

## Discussion

This study used fUS to examine visually evoked activity across the mouse visual system in a CLN3 disease model. By integrating functional responses with regional SCMAS accumulation, we identified distinct alterations along the two major visual pathways. In the geniculostriate pathway, *Cln3−/−* mice exhibited reduced activation in the anterior thalamus (aTH), whereas responses in V1 and posterior V2 remained largely comparable to WT controls. In contrast, regions along the extrageniculate pathway, including the midbrain, pTH, and aV2, showed increased activation in *Cln3−/−* mice. When considered together with SCMAS accumulation, these results highlight a pathway-dependent organization of pathology and functional responses across the visual system. Overall, SCMAS accumulation was more pronounced in the geniculostriate pathway than in the extrageniculate pathway, with the midbrain exhibiting the lowest level of pathology. These pathway-specific differences indicate that functional changes in the *Cln3−/−* visual system may be associated with non-uniform vulnerability across the visual network. Notably, regions along the extrageniculate pathway exhibited increased activation together with comparatively lower SCMAS accumulation.

One possible explanation for this increased engagement of the extrageniculate pathway is that the enhanced MB – pTH – aV2 fUS activation in Cln3−/− mice potentially reflects a compensatory recruitment of this pathway. When the primary geniculostriate pathway (retina – LGN – V1) is compromised, the extrageniculate pathway can exhibit adaptive functional compensation (Bridge et al., 2016; Nuzzi et al., 2018). Particularly, the SC – LP/pulvinar – extrastriatal cortex projection has been implicated in blindsight and residual vision, in which individual humans or non-human primates with V1 damage retain the capability to detect motion and salient stimuli without conscious perception (Celeghin et al., 2019; Leopold, 2012; Stoerig & Cowey, 1997). Increased responsiveness within this extrageniculate circuit has been proposed to represent a compensatory gain mechanism that preserves aspects of visual function despite primary thalamocortical dysfunction (Bridge et al., 2016; Kinoshita et al., 2019; Morris et al., 2001). Consistent with this interpretation, reduced fUS activation in aTH in *Cln3−/−* mice likely reflects an impaired primary thalamic response, whereas enhanced activation in MB, pTH, and aV2 suggests compensatory recruitment of extrageniculate pathways, potentially supporting V1 and pV2 activity via intracortical projections from aV2.

In lysosomal storage disorders, lysosomal pathology disrupts neuronal excitability by impairing mitochondrial function, increasing oxidative stress, and perturbing Ca^2+^homeostasis (Ballabio & Gieselmann, 2009; Plotegher & Duchen, 2017; Stepien et al., 2020). In the present study, SCMAS accumulation in the aTH, aV1, pV1, and pV2 was higher than that in the MB, pTH, and aV2 in Cln3−/− mice. These findings suggest that the extrageniculate pathway, which exhibited relatively lower pathological burden, may partially compensate for dysfunction within the geniculostriate pathway. Notably, the MB showed the lowest SCMAS accumulation among the examined visual regions, supporting the possibility of preferential recruitment of SC circuits and downstream pLP and visual cortical regions, analogous to blindsight-related networks that support residual visual processing (Kinoshita et al., 2019; Leopold, 2012). Together, these findings suggest that visual processing is reorganized at the circuit level in the presence of lysosomal storage pathology and identify potential targets for modulating compensatory network activity.

The experimental design in this study was chosen to balance imaging stability, physiological relevance, and minimal invasiveness. Imaging was performed transcranially with the skull left intact, avoiding craniotomy and skull thinning procedures. At the early ages examined in this study, the cranial bone is sufficiently thin to permit effective ultrasound transmission while minimizing surgical manipulation. Preserving the intact skull and suturing immediately after imaging facilitated rapid recovery and enabled animals to be reused for longitudinal studies or additional assessments (Li et al., 2017). In the present study, imaging was performed under light anesthesia rather than in awake head-fixed or freely moving conditions. While awake paradigms provide a more natural physiological state, they also introduce unavoidable movement-related activity (Brunner et al., 2021; Tiran et al., 2017). This consideration is particularly relevant in CLN3 disease, where photophobia has been reported and may involve both visual and somatosensory processing pathways (Maleki et al., 2012; Noseda et al., 2019). Body or facial movements during light stimulation could therefore activate overlapping somatosensory regions, complicating the interpretation of visually evoked responses. Maintaining controlled light anesthesia allowed us to prioritize stable, stimulus-driven activity within the visual system while minimizing motion-related confounds.

The anesthesia regimen was optimized based on pilot experiments and prior neuroimaging studies. Isoflurane concentrations around 1.5% produced pronounced vasodilation, elevating baseline cerebral blood volume (CBV) and reducing sensitivity to stimulus-evoked changes (Banoub et al., 2003; Iida et al., 1998). Although lowering the isoflurane concentration enhanced the relative CBV response, it also increased muscle-related motion artifacts. To balance vascular stability and motion suppression, dexmedetomidine was incorporated into the anesthesia protocol. This anesthesia combination, adapted from established resting-state fMRI protocols, partially counterbalances the vasodilatory effects of isoflurane with the vasoconstrictive effects of dexmedetomidine, improving hemodynamic stability (Grandjean et al., 2014). Moreover, low-dose isoflurane may suppress potential epileptic responses elicited by dexmedetomidine (Fukuda et al., 2013). Together, this approach maintained stable physiological conditions while preserving sensitivity to visually evoked CBV responses.

Another key methodological consideration was accounting for the temporal dynamics of neurovascular coupling by convolving the stimulus time course with a canonical hemodynamic response function (HRF) prior to correlation analysis. Although modeling the HRF within a General Linear Model (GLM) framework is standard practice in fMRI (Friston et al., 1998; Lindquist, 2008), it has been less frequently incorporated into fUS studies. Only a limited number of reports have applied HRF modeling in fUS activation mapping (Brunner et al., 2020; Brunner et al., 2021), and recent work has further characterized regional variability in HRF dynamics using fUS in awake mice (Erol et al., 2023). Supplementary comparisons (Supplementary Figure 1) of activation maps generated with and without HRF modeling illustrate the importance of accounting for the expected delay and temporal spread of the vascular response. Therefore, incorporating HRF modeling is critical for accurate detection of visually evoked activity, particularly under anesthesia, where the hemodynamic response shows delayed onset and extended duration.

Beyond the current findings, this fUS imaging framework provides a foundation for exploring photophobia and related circuit mechanisms in future studies. fMRI studies in migraine patients indicate an increased blood-oxygenation-level-dependent (BOLD) response in the SC, pulvinar, and visual and somatosensory cortices during light stimulation (Maleki et al., 2012; Panorgias et al., 2019). These findings suggest hypersensitization of subcortical and thalamocortical regions during photophobic episodes (Maleki et al., 2012; Panorgias et al., 2019). The enhanced visual fUS activation observed in the MB, pTH, and aV2 of *Cln3−/−* mice parallels findings reported in other disease conditions, suggesting abnormal amplification within the extrageniculate pathway that may contribute to photophobic phenotypes. Although photophobia has been reported in individuals with CLN3 disease (Kuper et al., 2021; Ouseph et al., 2016), the underlying circuit dysfunction remains poorly characterized. Most prior studies have focused on photoreceptor degeneration and retinal pigment epithelium atrophy associated with vision loss (Preising et al., 2017; Tang et al., 2021; Wright et al., 2020). Our findings reveal visually evoked central functional alterations, providing a foundation for investigating photophobic circuit dysfunction in mouse models of CLN3 disease.

The results in this study should also be considered in light of several limitations. First, fUS measures changes in cerebral blood flow rather than direct neuronal firing. As a hemodynamic proxy, its temporal dynamics are constrained by neurovascular coupling and thus do not reflect instantaneous neural activity. Second, although the spatial resolution of the probe used in this study (~100 *µ*m) was sufficient to resolve the ROIs at the regional scale, some nuclei, such as NOT and PPT, are relatively small structures in the mouse brain. Quantification in these areas is therefore more challenging because their small size increases sensitivity to minor registration misalignment, potentially contributing to variability in statistical estimates.

Despite these limitations, this study provides several methodological and conceptual advances for the understanding of CLN3 disease. The use of fUS enabled the investigation of brain-wide visual responses at early disease stages with high spatial resolution and sensitivity to hemodynamic changes, allowing simultaneous measurement of activity in both cortical and subcortical regions. Our results demonstrated that fUS can detect functional alterations across central visual pathways in CLN3 disease beyond retinal degeneration and ocular pathology. By combining fUS measurements with quantitative SCMAS pathology, we further examined how functional changes relate to regional pathological burden across distinct components of the visual system. Together, this work provides insight into early circuit-level dysfunction in CLN3 disease and identifies pathway-specific network alterations associated with regional pathological accumulation.

## Conclusion

This study demonstrates pathway-specific functional alterations in the visual system of young *Cln3−/−* mice using fUS imaging. We observed enhanced activation along the extrageniculate pathway together with maintained or reduced responses within the geniculostriate pathway. Combined with SCMAS quantification revealing region-dependent pathological burden, these findings suggest that pathway-level reorganization, especially involving midbrain-associated circuits, contributes to early functional alterations in CLN3 disease. These results further indicate non-uniform disease progression across central visual pathways and highlight the utility of fUS imaging for probing early dysfunction in neurodegenerative models.

## Data availability

The data and code supporting the findings of this study are currently available from the corresponding author upon reasonable request.

## CRediT authorship contribution statement

**Fanchao Yin**: Writing – Original Draft, Writing – Review & Editing, Conceptualization, Methodology, Software, Formal analysis, Investigation, Visualization, Data curation. **Yanya Ding**: Writing – Original Draft, Writing – Review & Editing, Conceptualization, Methodology, Software, Formal analysis, Investigation, Visualization, Data curation. **Hayley Chang**: Methodology, Resources. **Viollandi Prifti**: Methodology, Resources. **Jingyu Feng**: Methodology, Resources. **Edward G Freedman**: Resources, Funding acquisition. **John J Foxe**: Resources, Funding acquisition. **Marvin Doyley**: Writing – Review & Editing, Conceptualization, Supervision, Funding acquisition, Resources. **Kuan Hong Wang**: Writing – Review & Editing, Conceptualization, Supervision, Funding acquisition, Resources.

## Acknowledgements

This work was funded by the National Institutes of Health (P50 HD103536-7954, R21EB033122).

We thank members of Frederick J. and Marion A. Schindler Cognitive Neurophysiology Lab and Wang Lab for critical discussions. We thank Wentao Hu for valuable assistance with functional ultrasound imaging and helpful discussions.

We acknowledge the use of the Center for Musculoskeletal Research (CMSR) and the Center for Advanced Microscopy & Nanoscopy (CALMN) at the University of Rochester Medical Center for imaging service and support. These facilities are supported by the National Institutes of Health (AR069655, P50 HD103536).

