## Supplementary material for "Functional ultrasound imaging reveals pathway-specific visual system reorganization in young *Cln3*−/− mice"

**Supplementary Table 1. Summary of mixed-effects models for two pathways and fUS activation metrics.**

| Pathway | Metric | Analysis | Comparison/Effect | Test statistic (F) | Adjusted p-value |
| --- | --- | --- | --- | --- | --- |
| Extrageniculate | % Activation | Generalized linear mixed-effects model | ROI* | F(4,105) = 5775.5 | < 0.0001 |
|  |  |  | Genotype* | F(1, 105) = 4.8610 | 0.0030 |
|  |  |  | ROI × Genotype* | F(4, 105) = 1796.7 | < 0.0001 |
|  |  | Tukey’s multiple comparisons between WT and *Cln3-/-* | aMB |  | 0.0727 |
|  |  |  | pMB |  | 0.3484 |
|  |  |  | pTH* |  | 0.0497 |
|  |  |  | aPM* |  | 0.0356 |
|  |  |  | AL* |  | 0.0460 |
|  | Mean Z-score | Mixed-effects model | ROI* | F(2.091, 43.90) = 7.1040 | 0.0019 |
|  |  |  | Genotype | F(1, 21) = 2.2150 | 0.1515 |
|  |  |  | ROI × Genotype | F(2.091, 43.90) = 0.1586 | 0.8625 |
|  |  | Tukey’s multiple comparisons between WT and *Cln3-/-* | aMB |  | 0.3986 |
|  |  |  | pMB |  | 0.4546 |
|  |  |  | pTH |  | 0.1216 |
|  |  |  | aPM |  | 0.4051 |
|  |  |  | AL |  | 0.2538 |
| Geniculostriate | % Activation | Generalized linear mixed-effects model | ROI* | F(4,100) = 3070 | < 0.0001 |
|  |  |  | Genotype | F(1, 100) = 0.25171 | 0.6170 |
|  |  |  | ROI × Genotype* | F(4, 100) = 2542.1 | < 0.0001 |
|  |  | Tukey’s multiple comparisons between WT and *Cln3-/-* | aV1 |  | 0.2398 |
|  |  |  | pV1 |  | 0.8224 |
|  |  |  | pPM |  | 0.8224 |
|  |  |  | L |  | 0.8986 |
|  |  |  | aTH |  | 0.4375 |
|  | Mean Z-score | Mixed-effects model | ROI | F(2.543, 50.23) = 0.9871 | 0.3960 |
|  |  |  | Genotype | F(1, 21) = 0.0524 | 0.8212 |
|  |  |  | ROI × Genotype | F(2.543, 50.23) = 0.8269 | 0.4683 |
|  |  | Tukey’s multiple comparisons between WT and *Cln3-/-* | aV1 |  | 0.4586 |
|  |  |  | pV1 |  | 0.6775 |
|  |  |  | pPM |  | 0.8746 |
|  |  |  | L |  | 0.6795 |
|  |  |  | aTH* |  | 0.0321 |

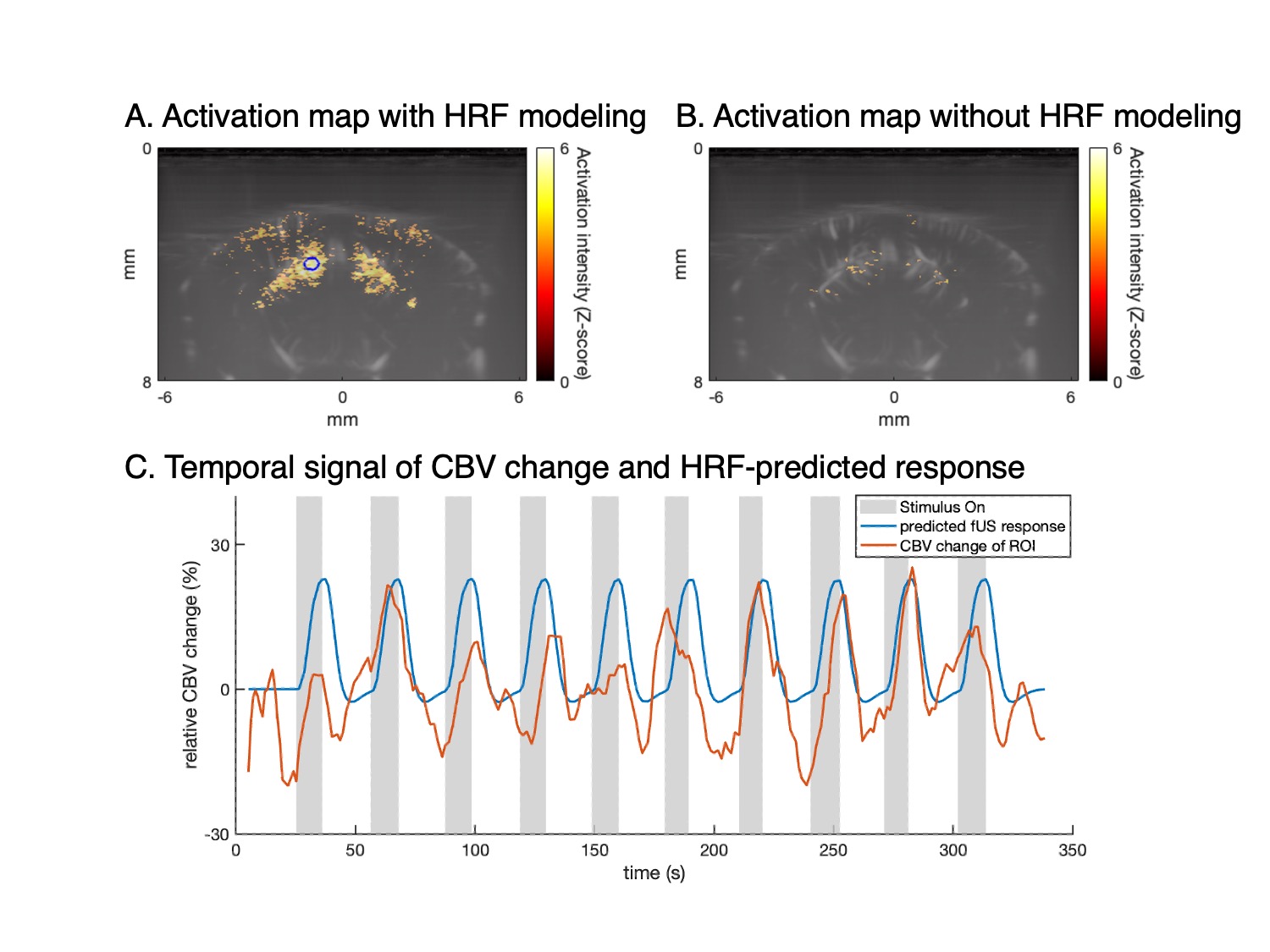

**Supplementary Figure S1. Effect of hemodynamic response modeling on activation detection and temporal dynamics.** Activation maps were generated either by correlating the fUS signal with the stimulus time course convolved with a canonical hemodynamic response function (HRF) (A) or by directly correlating the fUS signal with the stimulus time course without HRF modeling (B). The color scale represents activation intensity (Z-score). Incorporating HRF modeling yields stronger, more spatially coherent activation than direct correlation. The blue circle in panel (A) indicates the region of interest (ROI) used for temporal analysis. (C) The blue trace represents the predicted fUS response obtained by convolving the stimulus paradigm with the canonical HRF, and the red trace shows the measured relative cerebral blood volume (CBV) change extracted from the circled ROI. Gray shaded regions indicate stimulus ON periods, and white intervals indicate stimulus OFF periods. Accounting for the temporal characteristics of the hemodynamic response improves the alignment between the predicted response and measured CBV fluctuations.
